# Isometric handgrip contraction increases tibialis anterior intrinsic motoneuron excitability in a dose-dependent manner

**DOI:** 10.1101/2025.01.16.633442

**Authors:** Lucas Ugliara, Lucas B. R. Orssatto, Amilton Vieria, Gabriel S. Trajano

## Abstract

Persistent inward currents (PICs) contribution to motoneuron firing in the lower limb typically increase after a remote handgrip contraction, believed to result from diffuse increases of serotonergic input on the spinal cord. We investigated whether handgrip contraction intensity, duration, and/or impulse would affect tibialis anterior estimates of PICs. Multi-channel electromyograms were recorded from the tibialis anterior of 21 participants (18-40 years), during dorsiflexions at 20% of individual’s maximal torque, before and after four handgrip conditions: i) *80%15s*, 80% of their maximal handgrip strength sustained for 15s; ii) *40%15s*, 40% sustained for 15s; iii) *40%30s*, 40% sustained for 30s; and iv) *Control* (no handgrip). PICs contribution to self-sustained motoneuron firing was estimated with the delta frequency (ΔF) using the paired motor unit analysis. The ‘brace height’, normalised as a percentage of a right triangle (%rTri), was used to quantify the effects of PICs on the non-linearity of firing patterns, representing the neuromodulatory drive (metabotropic regulation of motoneuron excitability) onto the motoneurons. ΔF increased by 0.33 pulses per second (pps; 95%CI 0.16–0.49, *d*=0.47) after *40%30s* and by 0.24 pps (0.09–0.38, *d*=0.34) after *80%15s* but remained unchanged after *40%15s* and *Control*. Similarly, brace height increased by 2.24 %rTri (0.18–4.30, *d*=0.20) after *40%30s* and by 2.45 %rTri (0.64–4.25, *d*=0.22) after *80%15s;* remaining unchanged after *40%15s* and *Control*. The increase in PICs contribution to motoneuron firing induced by a remote handgrip contraction is impulse-dependent rather than intensity or duration. The parallel increases in ΔF and brace height suggest augmented neuromodulatory input onto the spinal cord.

## Introduction

Motoneurons rely on a complex control system for excitability, with neuromodulation as a key component. Unlike ionotropic systems, which directly convert synaptic inputs into action potentials via ion channel opening, neuromodulation operates through neurotransmitters that bind to specific receptors, initiating intracellular signalling pathways that regulate motoneuron responsiveness (Heckman et al., 2009). This neuromodulatory influence facilitates the generation of strong persistent inward currents (PICs) in spinal motoneurons, which increase cell excitability, accelerating, amplifying, and prolonging their discharge output activity for a given excitatory input (Heckman, Johnson, et al., 2008). This input–output gain mechanism can be adjusted based on the level of serotonin (5HT) and noradrenaline (NE) released from the raphe nuclei and locus coeruleus, respectively, which travel through the monoaminergic projections in the spinal cord (Heckman, Hyngstrom, et al., 2008), and activate their respective receptors located largely on the motoneurons’ dendrites. This activation promote an intracellular response via second messengers, leading to the opening of L-type Ca^2+^ and Na^+^ channels and facilitating PICs in a dose-dependent manner (Johnson & Heckman, 2014). It has been suggested that the level of neuromodulatory input onto the motoneurons could be adjusted according to the physical tasks demand (Heckman et al., 2009). Lower levels of monoaminergic drive would reduce PICs magnitude, allowing the motor system to perform delicate muscular actions with accuracy (e.g., threading a needle). Alternatively, moderate-to-maximal motor tasks (e.g., lifting heavy weights) would demand higher monoaminergic drive and an increased state of PIC facilitation (Johnson & Heckman, 2014). This theory is supported by the observation that the firing rate of neurons in the caudal raphe nuclei is influenced by the intensity of motor outflow, such that 5HT release is proportional to locomotion demands (Fenstermacher et al., 2024; Veasey et al., 1995). Importantly, much of current knowledge about the effects of neuromodulation on PICs and the role of 5HT in motoneuron gain control comes from invasive animal experiments and computational simulations (Heckman et al., 2009; Hounsgaard et al., 1988; Lee & Heckman, 2000). However, translating these findings to humans can be challenging, requiring non-invasive techniques.

In humans, researchers have used different strategies to investigate the contribution of 5HT and NE on PICs neuromodulation assessed with indirect measures. Pharmacological trials involving drugs that can change the concentration of 5HT and NE at the synaptic cleft have demonstrated their influence on estimates of PICs and different variables linked to motoneuron excitability (e.g., discharge rate, recruitment threshold) (D’Amico et al., 2013; Goodlich et al., 2023, 2024; Udina et al., 2010). Additionally, some studies have used muscle contraction of different intensities to theoretically increase monoaminergic input onto the spinal cord to understand its effects on PICs neuromodulation. These studies show that PICs contribution to motoneuron firing increase with contraction intensity levels of the same muscle group (Mackay et al., 2023; Orssatto et al., 2021). Other studies used remote handgrip contractions to, theoretically, increase 5HT availability via diffused serotonergic projections on the spinal cord and; thus, increase estimates of PICs on lower limb muscles (Mackay Phillips et al., 2023; Orssatto et al., 2022). This paradigm was supported by Wei et al. (2014), who showed that remote leg contractions increased force variance during a precision task with the palm or index finger. This effect was interpreted as 5HT-mediated, as force variation decreased in the presence of selective 5HT receptor blockers and reuptake inhibitors. These findings aligns with evidence that the firing rate of neurons in the caudal raphe nuclei and locus coeruleus is influenced, respectively, by the intensity of motor outflow and arousal state, such that 5HT and NE release is proportional to motor task or locomotion demands (Fenstermacher et al., 2024; Heckman, Johnson, et al., 2008; Veasey et al., 1995). Collectively, these findings align with evidence from animal experiments and support the assumption that it is feasible to investigate indirectly the contribution of serotonin, using contraction of remote muscle groups, to promote changes in motoneuron firing pattern in humans.

After establishing the efficacy of using remote handgrip tasks to increase 5HT input onto the spinal cord and facilitate tibialis anterior and soleus PICs (Mackay Phillips et al., 2023; Orssatto et al., 2022), the logical next step is to understand whether the PICs response could be differently influenced by mechanical aspects of the remote contraction (i.e., force intensity, duration, and impulse). This rationale is supported by evidence that higher contraction intensities increase estimates of PIC contribution to motoneuron self-sustained firing (e.g., ΔF) in muscles like the soleus, gastrocnemius (Orssatto et al., 2021) and tibialis anterior (Mackay et al., 2023), suggesting that intensity modulates PIC activation. Additionally, serotonergic neuron activity in the raphe nuclei of cats rises and maintains during prolonged locomotion until fatigue (*i.e,* the inability of the cat to maintain pace) (Jacobs et al., 2002) implying that sustained motor activity may also enhance PIC levels. To investigate this phenomenon, the present study used multi-channel electromyography to assess MU discharge rates, and the paired MU technique to estimate PICs contribution to motoneuron firing (Mesquita et al., 2024) following handgrip contractions at different intensity levels and contraction durations. In addition, we estimated the non-linearity caused by the effect of monoaminergic drive onto motoneuron discharge patterns using a quasi-geometric approach to calculate the ‘brace height’ of the ascending phase of the MU firing rates (Beauchamp et al., 2023). Given the evidence of the importance of the intensity of both muscle contraction for the estimates of PICs of motoneurons (Mackay et al., 2023; Orssatto et al., 2021) and greater levels of motor output inducing greater concentration of 5HT in the spinal cord (Jacobs et al., 2002), potentially leading to increasing motoneuron excitability, we hypothesize that the combination of both greater intensity and duration (i.e., impulse) of remote contraction will induce higher estimates of PIC activity.

## Methods

### Participants and ethical procedures

Twenty-three young adults participated in this study (Table 1). Inclusion criteria required volunteers to be within the age range of 18–40 years, have no history of musculoskeletal injuries in the tested limbs, and not be using medications that could affect the monoaminergic system (e.g., beta-blockers and selective serotonin reuptake inhibitors) (Thorstensen et al., 2024). Furthermore, participants were instructed to abstain from strenuous physical activities and alcohol consumption for 48 hours before the testing session. The Queensland University of Technology Human Research Ethics Committee approved the study (Reference number: 6770), and all participants signed a written informed consent prior to their participation.

**Table 1.** Participant characteristics.

| Participant characteristics |  |
| --- | --- |
| Age (years) | 31 (29, 33) |
| Sex (n) |  |
| Male | 15 |
| Female | 6 |
| Body mass (kg) | 79 (73, 86) |
| Height (cm) | 177(173, 181) |
| BMI (kg/m<sup>2</sup>) | 25 (24, 27) |
| Handgrip peak force (N) | 437 (386, 488) |
| Dorsiflexion peak torque (N·m) | 54 (49, 60) |

Participant characteristics data are presented as means with 95% confidence interval.

### Study design and testing procedures

The dorsiflexion and handgrip contractions were performed using the “preferred leg for kicking a ball” and the corresponding ipsilateral hand, respectively. For the dorsiflexion tasks, participants were positioned upright in an isokinetic dynamometer (Biodex System 4, Biodex Medical system, Shirley, NY) with the knee fully extended (0°), seat reclined at 70° of hip flexion and ankle in anatomical position. For the handgrip tasks, a handgrip dynamometer (model MLT004/ST, ADinstruments, Australia) was held with shoulder in the anatomic position at 0° flexion/extension, 0° abduction/adduction, and 10-15° external rotation, with the elbow flexed at 90°, while seated on the isokinetic dynamometer. A warm-up was performed alternating between handgrip and ankle dorsiflexion tasks, and consisted of progressive contraction levels (20%, 40%, 60%, 80% of perceived maximal effort) for ∼5 s each. Two minutes after, they performed two 4-s handgrip maximal voluntary contractions, with 60 s of interval between attempts. They were given two minutes of resting and then also performed two 4-s dorsiflexor maximal voluntary contractions, with 60 s of interval between attempts. Peak handgrip force and dorsiflexion torque were considered as the maximum value achieved during the maximal voluntary contractions. Thereafter, participants were familiarized with triangular-shaped ramped contractions to 20% of their dorsiflexion peak torque at 2%/s rate of torque increase and decrease (10-s up and 10-s down). Visual feedback was provided through a 23” computer monitor, guiding the participants to closely follow the real-time torque trajectory. This relative torque level was chosen for based on previous studies investigating the effects of a remote handgrip contraction on lower limbs ΔF (Mackay Phillips et al., 2023; Orssatto et al., 2022).

Five minutes after familiarization, four clusters of two sets of triangular-shaped contractions, interspersed with either a resting control condition or three distinct handgrip contractions were performed in a randomized order. During the control condition, the participants were asked to remain quiet and relaxed, avoiding to move/contract any muscle for 60 s in between the two triangular-shaped contractions. For the three conditions involving a remote handgrip contraction, participants were requested to quickly reach a relative grip force level and sustain it for a given time; the three conditions were i) handgrip contraction at 40% of their peak force sustained for 15 s (i.e., 40%15s); ii) handgrip contraction at 40% of their peak force sustained for 30 s (i.e., 40%30s), and iii) handgrip contraction at 80% of their peak force sustained for 15 s (i.e., 80%15s). Conditions 80%15s and 40%30s were impulse matched, 80%15s and 40%15s were time-matched, and 40%30s and 40%15s were intensity matched. This approach was adopted to allow us to investigate whether the increases in ΔF are affected by remote contraction time (40% sustained for 15 s vs 30 s), intensity (40% vs 80% sustained for 15 s), or impulse (40% sustained for 30 s and 80% sustained for 15 s vs 40% sustained for 15 s). The time between the two triangular-shaped contractions, performed before and after the control or handgrip conditions, was standardized at 60 s. Thus, a 30-s waiting period preceded the handgrip contraction lasting 30 s, while a 45-s waiting period preceded the handgrip contraction lasting 15 s. The second triangular-shaped contractions were performed immediately after the handgrip conditions. A 5-min rest interval was adopted between conditions. Figure 1 illustrates the study protocol design. In case abrupt/steep increases or decreases (>5% of peak torque) of torque were observed during the ascending or descending phase of the triangular-shaped contractions, the whole trial for the specific condition was excluded and repeated after 5 minutes. The participants were given only one extra attempt for each condition, when necessary. No further attempts were given to avoid the presence of fatigue, which can affect the outcomes obtained from the triangular contractions (Mackay et al., 2023). If the participant was not able to maintain consistency in the before and after triangular ramp-shaped contractions (i.e., avoid abrupt/steep increases or decreases in torque) for both attempts in a given condition, they were excluded from the analysis for that condition. Finally, the impulse (area under the force curve) performed at the handgrip conditions was calculated using the trapezoidal rule in LabChart software (version 7.3, ADinstruments, Australia).

**Figure 1.**
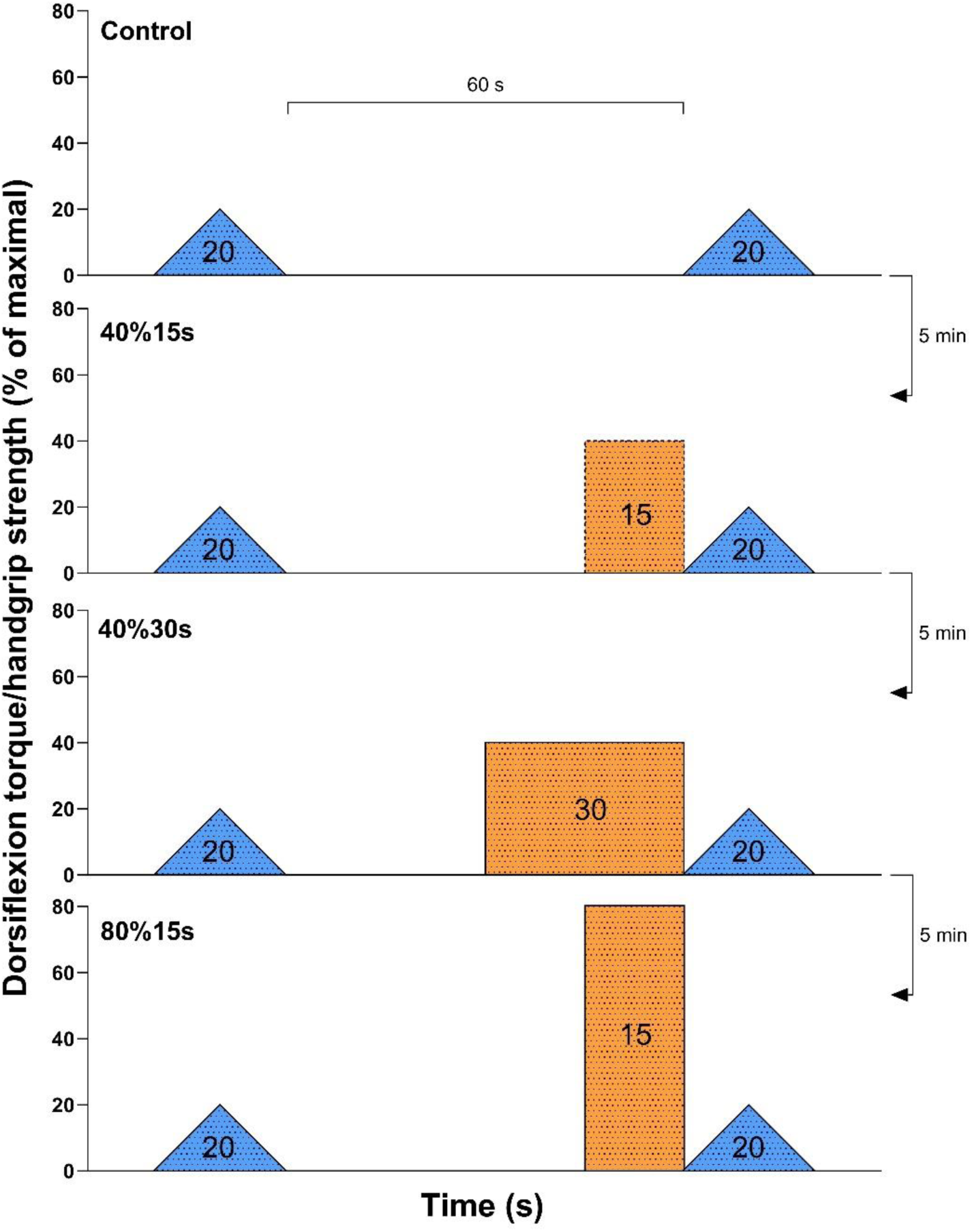
Study testing protocol. Blue triangles represent the triangular ramp-shaped dorsiflexion contraction with the time between these two tasks set at 60 s for all conditions; the dashed orange rectangle represents half handgrip impulse as the other two solid-lined orange rectangles; conditions were performed randomly with 5 min rest interval.

**Figure 2.**
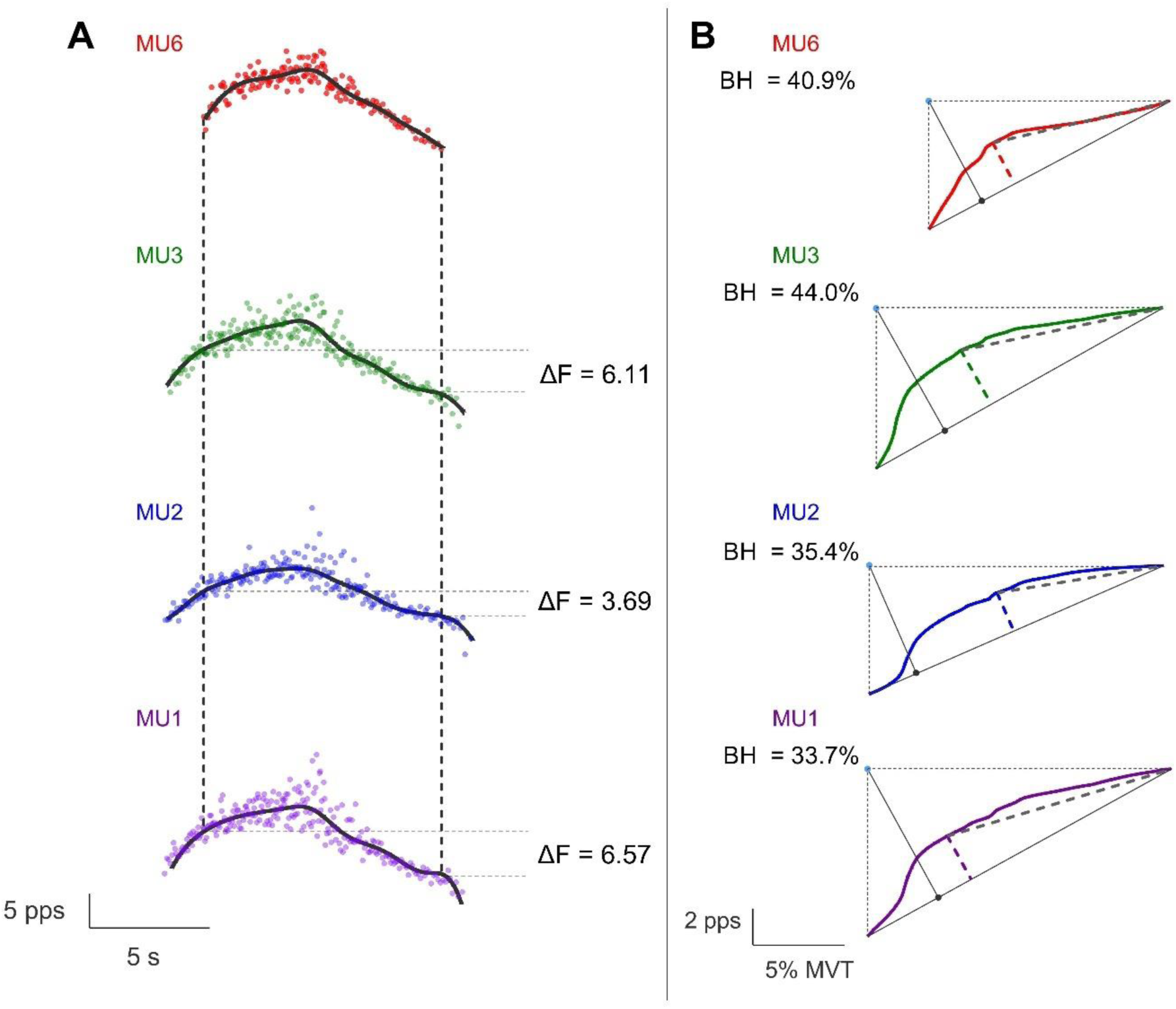
A, delta frequency (ΔF) calculation from a single participant for triangular-shaped contraction to 20% of maximal their maximal peak torque. *Red was used to represent firing of the test unit (top), paired with other three control units (green, blue and purple). Solid black lines, SVR smoothed curve of the MUs; dashed vertical black lines, moment of recruitment and decruitment of the test unit; dashed horizontal grey lines, firing of control units at the moment of recruitment and derecruitment of the test unit, used to calculate ΔF*. B, brace height and attenuation slope calculation for the same MUs in A. *Solid coloured lines, SVR smoothed curve of the MUs; dashed coloured lines, brace height; solid black line, orthogonal distance originating from the hypotenuse (blue dot) to the right triangle (black dot) formed by the moment of recruitment to derecruitment of each MU (dashed black lines); dashed grey lines, attenuation slope*.

### Multi-channel Electromyography Recordings and Analyses

The tibialis anterior skin area was prepared by shaving and cleansing with 70% isopropyl alcohol. A semi-disposable 64-channel electrode grid with an interelectrode distance of 8 mm (GR08MM1305, OT Bioelettronica, Turin, Italy), attached to a bi-adhesive foam layer and coated with a conductive paste (Ten20, Weaver and Company, Aurora, CO, USA) was positioned over the most prominent part of the tibialis anterior. The grounding electrode comprised dampened strap (WS2, OTBioelettronica, Torino, Italy) placed around the ankle joint.

Multi-channel electromyograms were recorded during the triangular ramp-shaped contractions using monopolar mode, amplified (256×), subjected to band-pass filtering (10–500 Hz), and digitally sampled at 2048 Hz, with a 16-bit wireless amplifier (Sessantaquattro, OT Bioelettronica, Turin, Italy), interfacing with OTBioLab + software (version 1.3.0., OT Bioelettronica, Turin, Italy) and were stored for subsequent offline analysis. The recorded data was processed offline using the DEMUSE software (Holobar & Zazula, 2007). The signals were band-pass filtered (20–500 Hz) through a second-order, zero-lag Butterworth filter. Subsequently, MU decomposition was performed using the convolutive kernel compensation (CKC) method (Holobar et al., 2014; Holobar & Zazula, 2007). Following decomposition, the same MUs were tracked across the before and after handgrip or control conditions, but not across conditions. The files for each contraction were concatenated and separation vectors (i.e., MU filters) were used to identify and generate the MU spike trains across the two contractions (Del Vecchio et al., 2019; Holobar et al., 2014; Holobar & Zazula, 2007). Then, a trained investigator (LU) examined the MU spike trains, and, when necessary, edited the discharge patterns of the MUs (Del Vecchio et al., 2020). Only MUs with a pulse-to-noise ratio ≥ 30 dB were retained for presenting higher reliability (sensitivity > 90% and false alarm rates < 2%) (Holobar et al., 2014). The edited MUs were quality-checked by a researcher, blinded for each condition, (LBRO) with extensive experience in decomposing, tracking and editing of MUs. When no MU was identified in both contractions for a given condition, the participant was not included in the respective condition analysis.

#### Estimation of PIC contribution to motoneuron self-sustained firing (ΔF) and peak discharge rate

The MU discharge events were converted into instantaneous discharge rates and smoothed with support vector regression machine learning fit (Beauchamp et al., 2022). PIC contribution to motoneuron firing was estimated using the paired MU analysis (Afsharipour et al., 2020). MUs with lower recruitment thresholds (control units) were paired with units of higher recruitment threshold (test units). ΔF was calculated as the change in discharge rates of the control MU from the onset of recruitment to the point of de-recruitment of the test unit (i.e., discharging hysteresis) (Afsharipour et al., 2020). MUs were paired when the following criteria was obtained between control and test units i) a rate-to-rate correlation threshold of r ≥ 0.7, ii) test – control unit recruitment time >1 s, iii) the control unit’s discharge rate at test unit recruitment minus the control unit’s peak discharge rate was > 0.5 pps, iv) control unit derecruitment time > test unit derecruitment time (Afsharipour et al., 2020; Hassan et al., 2020; Vandenberk & Kalmar, 2014). ΔFs calculated for each individual test unit were averaged across control units to yield a singular ΔF value for each corresponding test MU. In the case that no MU pair was identified in both contractions for a given condition, the participant was not included in the analysis for that respective condition(s). The highest value derived from the support vector regression fit curve was determined as peak discharge rate. The relative torque (%) produced during the recruitment time of each MU was identified as MU recruitment threshold.

#### Quantification of neuromodulatory and inhibitory drive in MUs firing changes

We used the quasi-geometric approach to quantify the neuromodulatory and inhibitory drive onto the MUs, as proposed by Beauchamp et al. (2023). Firstly, we generated a straight line (hypothenuse) from the discharge rate at MU recruitment to the peak discharge rate of the support vector regression (SVR) smoothed curve. Then, we calculated the ‘brace height’, which represents the maximum deviation of the smoothed discharge rates trace from linearity (hypothenuse). The maximal orthogonal distance between the hypothenuse and the smoothed MU discharge trace was considered the brace height. Thereafter, brace height was normalized as a percentage of the maximal orthogonal distance between the straight line and a right triangle in which sides originate from the hypotenuse. We also calculated the ‘attenuation slope’, which is associated with the inhibitory input effect on MU discharge (Beauchamp et al., 2023). Attenuation slope was calculated from the brace height insertion on the smoothed discharge rates to the peak discharge rates of the ascending phase.

## Data and statistical analyses

Linear-mixed effects models were investigated utilizing the *robustlmm* package (Kuznetsova et al., 2017). We estimated marginal mean differences between handgrip conditions in ΔF, peak discharge rate, brace height and attenuation slope along with 90% and 95% confidence intervals (CI), using the emmeans package (Lenth et al., 2023). Each MU was treated as repeated measure and nested for each participant, including a random intercept for each participant to consider for the correlation between repeated observations on each individual (i.e., 1| participant/MU ID). For ΔF, four distinct models were fitted: 1) time and condition were included as fixed effects and a random intercept and slope (ΔF) for each participant; 2) similar to model 1, but included peak discharge rate as covariate; 3) similar to model 1, but included recruitment threshold as covariate; 4) similar to model 1 but included both peak discharge and recruitment threshold as covariates. All the four models resulted in similar outcomes; thus we presented the findings derived from Model 4, which exhibited superior fit (as indicated by the lowest Akaike’s information criteria and Bayesian information criteria). We performed an additional analysis by adding sex as fixed effect to ΔF to investigate a possible effect of sex on results, but no time by condition by sex interaction effect [β = 0.32 (−0.29, 0.92) pps, SE = 0.32; t = 0.31] and main effect of sex [β = –0.80 (−1.76, 0.16) pps, SE = 0.49; t = –1.64] were observed, so sex was removed from the model. For all other variables except by recruitment thresholds (i.e., peak discharge rate, brace height and attenuation slope), a model with time and condition as fixed effects and recruitment thresholds as covariate was employed. For recruitment thresholds, the model included time and condition as fixed effects. In addition, the mean impulse for each condition were compared using the *lmerTest* package for linear mixed effects models analysis. Pairwise *post hoc* analysis was used for significant effects observed. Cohen’s d effect sizes were computed for the differences between after and before handgrip conditions, leveraging the population standard deviation (σ) estimated from corresponding robust linear mixed-effects models as the denominator (i.e, d = mean difference / σ). For interpretation, we considered Cohen’s d <0.20 trivial; 0.20–0.49 small; 0.50–0.79 moderate; ≥0.80 large. A significance threshold of 5% (α level) was adhered to for all tests. All the analyses were conducted in a free software environment (RStudio, version 2024.12.1). The complete dataset and corresponding R script are accessible at https://github.com/lugliara/PICs.

## Results

### Participant characteristics and MU identification

One female participant was excluded from all the analyses due to inconsistencies in performing the ramp-shaped triangle contractions in all the conditions and another female participant was excluded because no MUs were identified at any condition. The final analysis included data from 21 participants. Among these 21 participants, all of them were included in the control and 80% of peak force for 15 s conditions (80%15s). For the 40% of peak force for 15 s (40%15s), data from five participants were excluded from the analyses because they were unable to consistently follow the force trace path during the triangular-shaped contractions without steep increases and decreases of force production (N = 3), or MUs did not meet essential prerequisites to form pairs (N = 2). For the 40% of peak force for 30 s (40%30s), data from three participants were excluded from the analyses because they were unable to follow the feedback path (i.e., detected abrupt/steep increases or decreases >5% of peak torque) during the triangular-shaped contractions (N = 2) or no MUs were detected (N = 1).

341, 249, 245 and 316 MUs which were matched before and after the control (mean (95% CI) = 16.2 (13.9, 18.6), N = 21), 40%15s (15.6 (13.3, 17.8), N = 16), 40%30s (13.6 (10.8, 16.4), N = 18) and 80%15s (15.0 (12.2, 17.9), N = 21) conditions, respectively. This resulted in 199 (9.5 (7.8, 11.2), N = 21), 146 (9.1 (7.2, 11.1), N = 16), 137 (7.6 (5.7, 9.6), N = 18) and 191 (9.1 (6.8, 11.4), N = 21) test units, respectively.

#### Handgrip contraction impulse

The impulse generated during the handgrip contractions during 40%30s (estimated marginal mean = 5639 (95%CI: 5054, 6224) N·s) and 80%15s (5667 (95%CI: 5082, 6252) N·s) were similar (p = 0.868), but superior than 40%15s (2933 (95%CI: 2348, 3519) N·s, p < 0.001).

### MU outcomes

#### ΔF

A time by condition interaction effect was observed β = –0.40 (−0.62, –0.18) pps, SE = 0.11; t = –3.60]. ΔF increased from before to after the intervention on 40%30s [0.33 (0.16, 0.49) pps, *d* = 0.47 (0.23, 0.72)] and 80%15s [0.24 (0.09, 0.38) pps, *d* = 0.34 (0.14, 0.55)], but remained unchanged on 40%15s [0.10 (−0.06, 0.26) pps, *d* = 0.15 (−0.09, 0.38)] and control [-0.07 (−0.21, 0.06) pps, *d* = –0.11 (−0.31, 0.09)] conditions (Figure 3).

**Figure 3.**
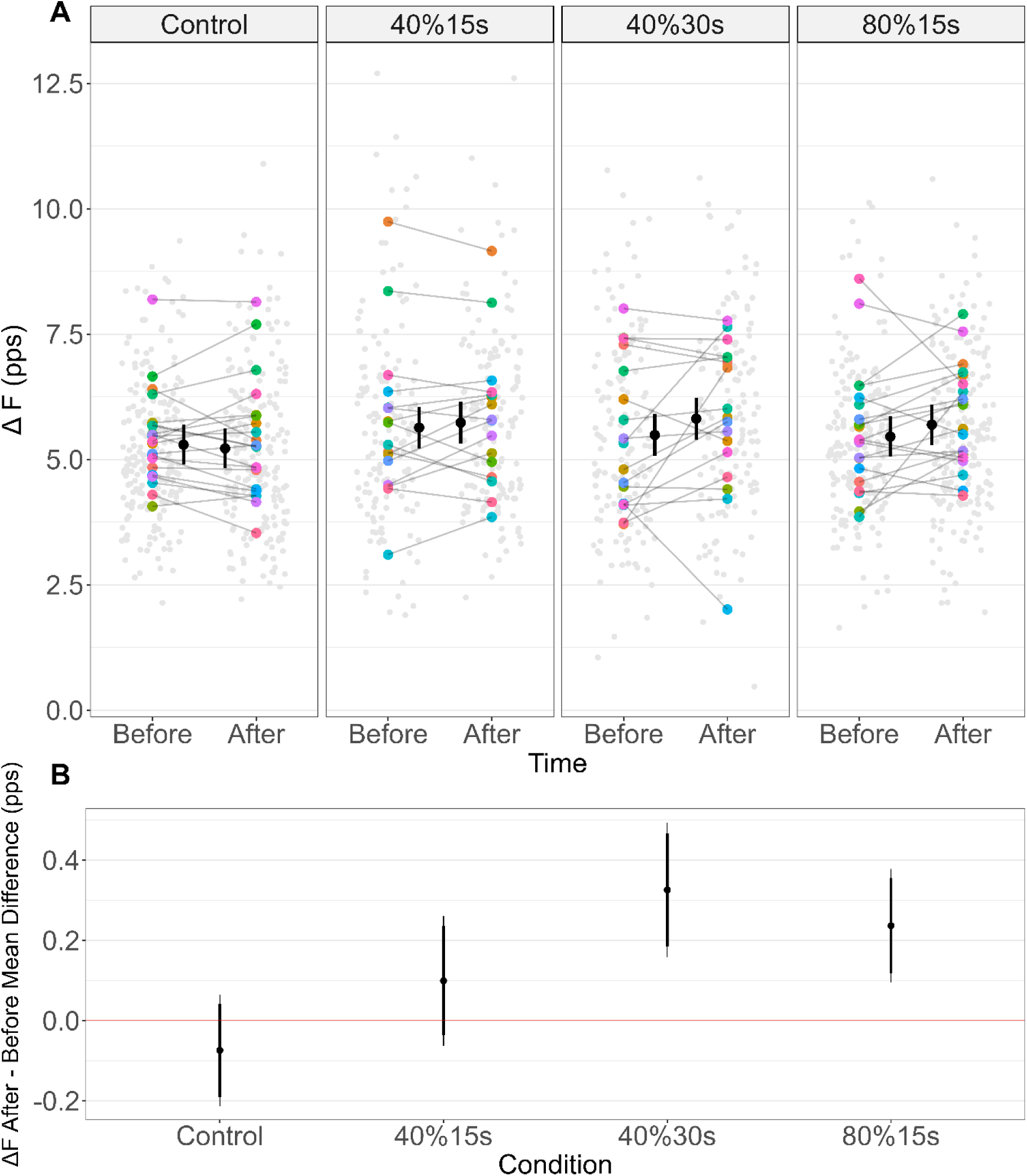
A, ΔF before and after the Control, handgrip to 40% of maximal isometric force for 15 s (40%15s) or 30 s (40%30s), and to 80% of maximal isometric force for 15 s (80%15s) conditions. *The estimated marginal means (black circles) and respective 95% confidence intervals are offset to the middle. Individual data points (averaged ΔF per participant) are coloured by participants and individual test units value are plotted in light grey. pps, pulses per second. ΔF remained unchanged before and after Control and 40%15s conditions but increased at 40%30s and 80%15s.* B, Estimated marginal mean differences (After – Before each condition) for ΔF in the control, handgrip at 40% of maximal isometric force for 15 s or 30 s, and at 80% of maximal isometric force for 15 s conditions. *Significant increases were observed on the handgrip at 40%30s and 80%15s conditions (not crossing the “zero” red line. The thick inner line and the thin outer line represent the 90% and 95% confidence intervals, respectively*.

**Figure 4.**
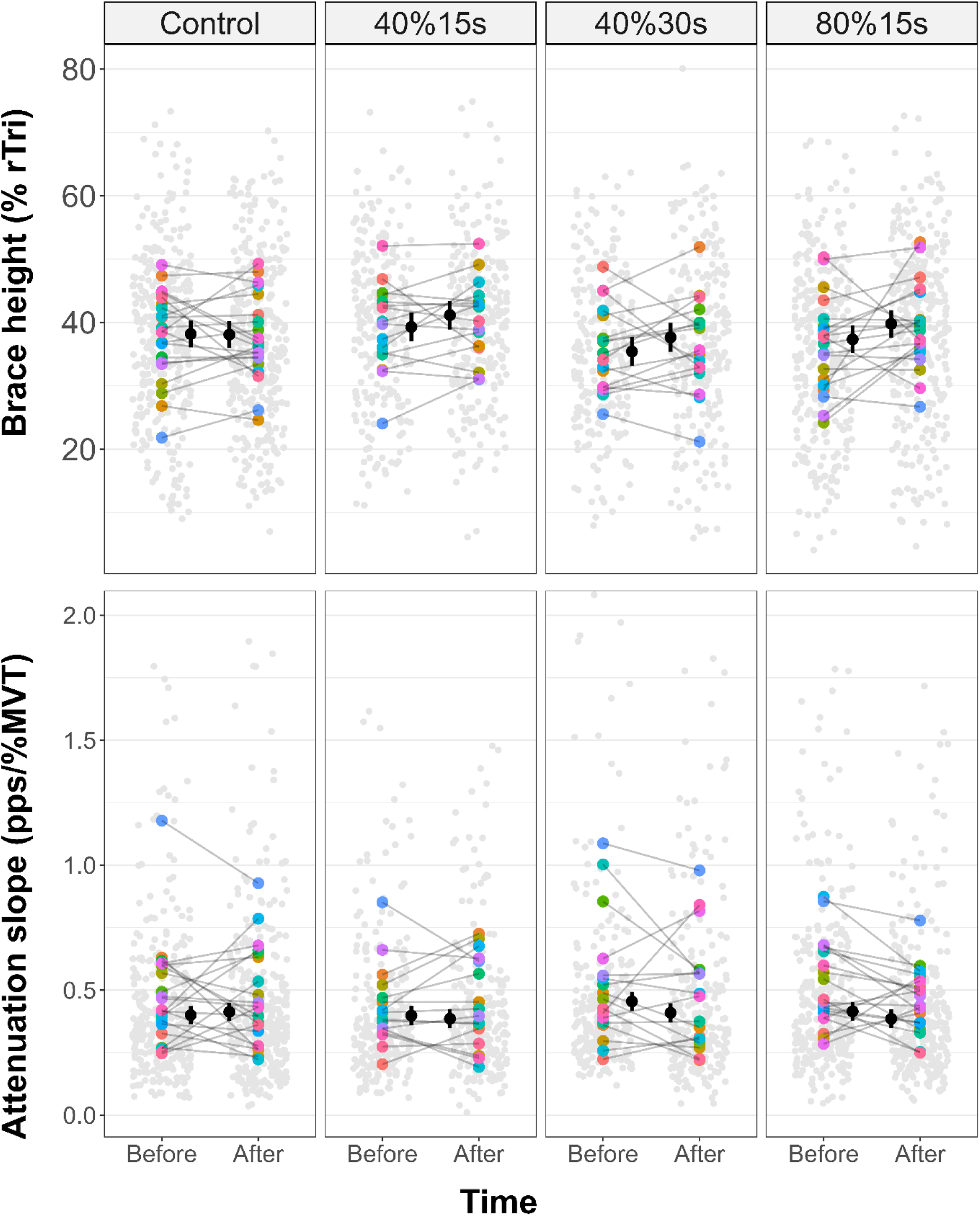
Top, brace height; bottom, attenuation slope; before and after the Control, handgrip to 40% of maximal isometric force for 15 s (40%15s) or 30 s (40%30s), and to 80% of maximal isometric force for 15 s (80%15s) conditions. *The estimated marginal means (black circles) and respective 95% confidence intervals are offset to the middle. Individual data points (averaged brace height and attenuation slope per participant) are coloured by participants and individual motor units value are plotted in light grey. pps, pulses per second; MVT, maximum voluntary torque; % rTri, percentage of the right triangle. Both brace height and attenuation slope remained unchanged before and after Control and 40%15s conditions but, respectively, increased and decreased at 40%30s and 80%15s. Note: the y-axis for attenuation slope has been limited to 2 pps/%MVT to enhance the visualization of data points and their 95% confidence intervals. A full-range version of the figure is available at* https://github.com/lugliara/PICs.

#### Brace height

A time by condition interaction effect was observed β = –2.56 (−5.06, –0.06) % rTri, SE = 1.27; t = –2.01]. Brace height increased from before to after the intervention on 40%30s [2.24 (0.18, 4.30) % rTri, *d* = 0.20 (0.02, 0.39)] and 80%15s [2.45 (0.64, 4.25) % rTri, *d* = 0.22 (0.06, 0.39)], but remained unchanged on 40%15s [1.86 (−0.15, 3.87) % rTri, *d* = 0.17 (−0.01, 0.35)] and control [ –0.11 (−1.84, 1.61) % rTri, *d* = –0.01 (−0.17, 0.15)] conditions.

#### Attenuation slope

A time by condition interaction effect was observed β = 0.06 (0.02, 0.10) pps/%MVT, SE = 0.02; t = 2.73]. Attenuation slope decreased from before to after the intervention on 40%30s [-0.05 (−0.08, –0.01) pps/%MVT, *d* = –0.27 (−0.46, –0.08)]and 80%15s [-0.03 (−0.06, –0.00) pps/%MVT, *d* = –0.17 (−0.34, –0.01)], but remained unchanged on 40%15s [-0.01 (−0.04, 0.02) pps/%MVT, *d* = –0.07 (−0.26, 0.11)] and control [0.01 (−0.01, 0.04) pps/%MVT, *d* = 0.07 (−0.08, 0.23)].

#### Peak discharge rates

A time by condition interaction was observed β = –0.50 (−0.62, –0.37) pps, SE = 0.06; t = –7.64]. Peak discharge rates decreased from before to after the intervention only on control [-0.29 (−0.38, –0.21) pps, *d* = –0.52 (−0.67, –0.36)] and increased on 80%15s [0.20 (0.11, 0.29) pps, *d* = 0.35 (0.19, 0.51)] condition, but remained unchanged on 40%15s [0.06 (−0.04, 0.16) pps, *d* = 0.10 (−0.08, 0.28)] and 40%30s [-0.09 (−0.19, 0.02) pps, *d* = –0.16 (−0.34, 0.03)].

#### Recruitment thresholds

A time by condition interaction effect was observed β = –0.89 (−1.17, –0.60) % of peak torque, SE = 0.15; t = –6.01]. Recruitment thresholds increased from before to after the intervention only on 40%15s [0.51 (0.29, 0.73) % of peak torque, *d* = 0.42 (0.24, 0.60)] and 40%30s [0.31 (0.09, 0.54) % of peak torque, *d* = 0.26 (0.07, 0.44)] conditions, and decreased on control [-0.38 (−0.57, –0.19) % of peak torque, *d* = –0.31 (−0.47, –0.16)] condition, but remained unchanged on 80%15s [-0.06 (−0.26, 0.14) % of peak torque, *d* = –0.05 (−0.21, 0.11)] condition.

Table 2 describes the before and after estimated marginal means and mean differences for each MU outcome and tested conditions.

**Table 2.** Estimated marginal means and mean differences (95% confidence interval lower and upper limits) for ΔF, peak discharge rates, recruitment thresholds, brace height, and attenuation slope for handgrip and control conditions.

| | $\Delta F$<br>(pps) | Brace height<br>(% rTri) | Attenuation slope<br>(pps/%MVT) | Peak discharge rate<br>(pps) | Recruitment thresholds<br>(% of peak torque) |
| --- | --- | --- | --- | --- | --- |
| **Control** |  |  |  |  |  |
| Before | 5.29 (4.90, 5.69) | 38.2 (36.1, 40.3) | 0.40 (0.36, 0.44) | 17.2 (16.1, 18.2) | 8.54 (7.16, 9.92) |
| After | 5.22 (4.82, 5.62) | 38.1 (36.0, 40.2) | 0.41 (0.38, 0.45) | 16.9 (15.8, 17.9) | 8.16 (6.79, 9.54) |
| After-before | -0.07 (-0.21, 0.06) | -0.11 (-1.84, 1.61) | 0.01 (-0.01, 0.04) | **-0.29 (-0.38, -0.21)** | **-0.38 (-0.57, -0.19)** |
| **40%15s** |  |  |  |  |  |
| Before | 5.64 (5.22, 6.05) | 39.3 (37.0, 41.6) | 0.40 (0.36, 0.44) | 17.1 (16.0, 18.1) | 8.08 (6.66, 9.49) |
| After | 5.73 (5.32, 6.15) | 41.2 (38.9, 43.4) | 0.39 (0.35, 0.42) | 17.1 (16.1, 18.2) | 8.58 (7.17, 10.00) |
| After-before | 0.10 (-0.06, 0.26) | 1.86 (-0.15, 3.87) | -0.01 (-0.04, 0.02) | 0.06 (-0.04, 0.16) | **0.51 (0.29, 0.73)** |
| **40%30s** |  |  |  |  |  |
| Before | 5.49 (5.07, 5.91) | 35.4 (33.1, 37.7) | 0.46 (0.42, 0.49) | 17.4 (16.3, 18.4) | 8.32 (6.91, 9.74) |
| After | 5.81 (5.39, 6.24) | 37.7 (35.4, 39.9) | 0.41 (0.37, 0.45) | 17.3 (16.2, 18.3) | 8.63 (7.22, 10.05) |
| After-before | **0.33 (0.16, 0.49)** | **2.24 (0.18, 4.30)** | **-0.05 (-0.08, -0.01)** | -0.09 (-0.19, 0.02) | **0.31 (0.09, 0.54)** |
| **80%15s** |  |  |  |  |  |
| Before | 5.46 (5.05, 5.86) | 37.3 (35.2, 39.5) | 0.412 (0.38, 0.45) | 17.4 (16.4, 18.5) | 9.00 (7.61, 10.40) |
| After | 5.69 (5.29, 6.10) | 39.8 (37.6, 42.0) | 0.39 (0.35, 0.42) | 17.6 (16.6, 18.7) | 8.94 (7.55, 10.33) |
| After-before | **0.24 (0.09, 0.38)** | **2.45 (0.64, 4.25)** | **-0.03 (-0.06, -0.00)** | **0.20 (0.11, 0.29)** | -0.06 (-0.26, 0.14) |

$\Delta F$ , $\Delta$ frequency; % rTri, percentage of the right triangle; MVT, maximum voluntary torque. Bold values indicate significant changes (95%CI not including zero).

## Discussion

We investigated whether some characteristics of the remote handgrip contraction, including duration (40%15s vs 40%30s), the intensity of effort (40%15s vs 80%15s), and the impulse (40%30s and 80%15s vs 40%15s) affect tibialis anterior PICs. Additionally, quasi-geometric analyses were used to estimate the contribution of neuromodulatory (i.e., brace height) and inhibitory (i.e., attenuation slope) inputs to PIC responses. The main findings of this study were: 1) ΔF increased after the matched higher handgrip impulse conditions; 2) brace height also increased under the same conditions; 3) attenuation slope decreased consistently after the higher handgrip impulse conditions. These findings suggest that the mechanical impulse of a remote contraction was the primary driver of increased tibialis anterior ΔF, likely due to enhanced neuromodulation and a shift in the inhibitory pattern.

The changes we observed in ΔF, brace height, and attenuation slope between the two highest handgrip impulse conditions (80%15s and 40%30s), compared to 40%15s and Control, support our hypothesis, indicating that a given impulse threshold might be necessary to influence increases in motoneuron recruitment-derecruitment hysteresis of tibialis anterior MUs. The small increase in ΔF after handgrip contractions at the 40%30s (d = 0.47) and 80%15s (d = 0.34) conditions were similar to the changes observed by Mackay et al., (2023) in soleus motoneurons, d = 0.30 and Orssatto et al., (2022) in tibialis anterior motoneurons, d = 0.55, both using a similar protocol to our 40%30s condition. Such changes on ΔF can be influenced by either neuromodulatory and/or inhibitory input onto the motoneurons (Beauchamp et al., 2023). The neuromodulatory input caused by the presence of monoamines is the main mechanism controlling PICs activity. In animals, voltage-clamp experiments provided strong evidence that PICs are led by the activation of voltage-gated L-type Ca^2+^ (Eckert & Lux, 1976; Svirskis & Hounsgaard, 1997) and Na^+^ (Harvey et al., 2006; Schwindt & Crill, 1977) channels. Subsequent studies showed the input of 5HT and NE activating specific G-protein coupled receptors at motoneurons’ dendrites and soma were responsible for triggering intracellular signalling cascade that modulate the properties of these ion voltage-gated channels (Harvey et al., 2006; Heckman et al., 2009; Perrier & Hounsgaard, 2003). In humans, oral administration of amphetamine, which is assumed to enhance the presynaptic release of NE, also increased estimates of ΔF (Udina et al., 2010), suggesting the neuromodulation mechanism observed in animals could also be present in humans. Moreover, Wei et al. (2014) demonstrated that 5HT modulates motoneurons input-output gain by using selective 5HT reuptake inhibitors and receptor blockers. More recent studies used multi-channel electromyography (EMG) during voluntary isometric ramp-shaped contractions and showed evidence of decreased MU excitability in presence of 5HT blockers by measuring ΔF and/or others estimates of PIC activity (Goodlich et al., 2023, 2024). Furthermore, studies found higher ΔF during ramped-shaped contractions at greater intensity (Mackay et al., 2023; Orssatto et al., 2021), suggesting higher voluntary muscle activity in humans is associated with higher PIC activity and, likely, higher 5HT and NE concentration into the spinal cord. Therefore, it is reasonable to assume that the increase in ΔF observed in this study after handgrip contractions with higher impulse could have been caused by a transient increase in monoamines concentration within the spinal cord; however, this mechanism was not directly tested in this study and future studies with drugs affecting monoamines concentration might be able to address this mechanism.

We used brace height to estimate neuromodulation contribution to motoneurons firing changes, since it was proposed to be influenced by neuromodulation, likely resulting from increased serotonergic and/or noradrenergic input (Beauchamp et al., 2023). Thus, the increases we observed in brace height at the 40%30s (*d* = 0.20) and 80%15s (*d* = 0.22) conditions theoretically suggest remote handgrip contraction induce greater neuromodulation. In addition, PICs are highly sensitive to inhibition (Hultborn et al., 2003; Kuo et al., 2003), an important feature to control unwanted movement that could occur as a consequence of neuromodulation by the diffuse descending projections of the monoaminergic system (Heckman et al., 2009). Previous studies observed inhibition responses at spinal level through different stimuli, such as small changes in antagonist muscles length (Hultborn et al., 2003; Hyngstrom et al., 2007), tendon vibration (Matthews, 1966; Orssatto et al., 2022; Pearcey et al., 2022), passive muscle stretching (Trajano et al., 2014), and voluntary co-contraction (Gomes et al., 2024), likely induced by activation of a disynaptic inhibitory circuit activated by muscle spindle primary Ia afferents (Crone et al., 1987; Kuffler et al., 1951). However, it is unlikely that changes in Ia inhibition would elicit inhibitory inputs in the different conditions used in the present study due to the consistency of the isometric ramp-shaped contraction we used before and after the remote contraction. Inhibition input might also originates at supraspinal or segmental neural mechanisms, like the observed by Renshaw cell activity (Hultborn & Pierrot-Deseilligny, 1979) and those generated by nociceptive inputs (Heckman et al., 2009). We used attenuation slope to estimate the influence of inhibitory input since the inhibition pattern was identified as a significant predictor of this measure (Beauchamp et al., 2023). The small and trivial decrease in attenuation slope at 40%30s (*d* = –0.27) and 80%15s (*d* = –0.17) might indicate a transition from a tonic inhibitory pattern to inhibitory commands that are more reciprocal to excitation (i.e., push-pull excitation-inhibition synaptic control) (Škarabot et al., 2023). However, caution is warranted before attributing this decrease solely to reduced inhibitory input, as ΔF sensitivity to inhibition patterns depends on neuromodulation levels (Beauchamp et al., 2023). Additionally, MUs exhibiting larger brace height would theoretically be associated with lower attenuation slope (Mesquita et al., 2024). Thus, our findings suggest that, once a certain level of impulse is reached, remote contractions performed at either lower or higher intensity/duration result in similar increase in the estimated contribution of PICs to motoneuron self-sustained firing (ΔF). This effect was likely driven by increased neuromodulation onto the spinal cord, as indicated by the small increase observed in brace height, while attenuation slope might reflect a shift in the pattern of inhibitory input.

Complementarily, we investigated peak discharge rates and recruitment threshold since both of which are associated with motoneuron excitability and might also be influenced by PIC activity (Mesquita et al., 2024; Orssatto et al., 2022). Moderate and small decreases in peak discharge rates (d = –0.52) and recruitment threshold (d = –0.38) were observed under the Control condition, contrasting with previous studies using a similar protocol, where no changes in Control were reported for tibialis anterior (Orssatto et al., 2022) or soleus motoneurons (Mackay Phillips et al., 2023). Unlike these studies, participants in our study performed three distinct handgrip tasks in a randomized order with 5-minute intervals, which may have mitigated task carryover effects on ΔF but not on peak discharge rates or recruitment threshold. Furthermore, we found a small increase (d = 0.35) in peak discharge rates only on 80%15s condition, whereas recruitment threshold increased in both the 40%15s (d = 0.42) and 40%30s (d = 0.26) conditions. Previous studies reported small increases in peak discharge rates under the 40%30s condition (Orssatto et al., (2022): d = 0.36; Mackay Phillips et al., (2023): d = 0.37), but while Orssatto et al. (2022) observed no changes in recruitment threshold, Mackay et al. (2023) did not report recruitment threshold results. These findings suggest that PICs may differentially modulate MU ΔF, peak discharge rates, and recruitment threshold, with these variables providing distinct insights into motoneuron excitability.

### Final considerations and future directions

This study combined robust non-invasive methods to estimate the influence of motor task on PIC activity in humans, likely via increased 5HT input to the spinal cord. Using an innovative EMG protocol allowed us to non-invasively assess key metrics of motoneuron excitability (e.g., ΔF, brace height and attenuation slope) without pharmacological intervention (Beauchamp et al., 2023). We also minimized the potential for type I error through rigorous statistical analysis (Yu et al., 2022). On the other hand, we assessed tibialis anterior MUs recruited during a ramp-shaped contraction to 20% of maximal force. Thus, caution should be taken when extrapolating our findings beyond lower-threshold units, at lower intensity contractions, and from the tibialis anterior muscle. Additionally, direct measurements of 5HT concentration in the spinal cord in vivo remain technically impossible, and we must still rely on indirect methods to infer its effects. Lastly, while we focused on 5HT and NE contributions to PICs, the potential influence of other neuromodulators should be considered in future studies. For example, evidence suggests that spinal interneurons may modulate motoneuron excitability via muscarinic m2 receptor activation, affecting resting K^+^ conductance (Miles et al., 2007).

Future studies could incorporate additional methods alongside EMG to clarify findings further. For instance: (1) given that both 40%30s and 80%15s conditions successfully increased ΔF in young adults, similar protocols could assess neuromodulatory capacity in populations with potentially reduced 5HT input or impaired PIC activation, such older adults, individuals with neurodegenerative diseases, or partial spinal cord injuries; (2) further investigations are desirable whether impulse-specific effects on PIC modulation differ across muscle groups, as variations in motoneuron distribution may affect responsiveness to remote contraction-induced neuromodulation;

## Conclusion

This study provides novel insights into the mechanisms through which remote voluntary muscle contractions influence motoneuron excitability, estimated by ΔF in tibialis anterior motor units. By manipulating both the intensity and duration of a handgrip task, we identified contraction impulse as a critical factor driving changes in motoneuron recruitment-derecruitment hysteresis, which likely reflects in increased PICs activity. Specifically, higher ΔF values in remote task conditions with matched contraction impulse suggest that both higher and lower intensity/duration conditions can lead to increased PIC modulation, provided that a sufficient impulse threshold is met. The increased brace height observed in the impulse-matched conditions support our hypothesis that increased tibialis anterior ΔF following handgrip remote contraction may result from increased neuromodulation, possibly by augmented serotonergic input onto the spinal cord.

